# Extending differential gene expression testing to handle genome aneuploidy in cancer

**DOI:** 10.1101/2025.03.29.646108

**Authors:** Katsiaryna Davydzenka, Giulio Caravagna, Guido Sanguinetti

## Abstract

Genome aneuploidy, characterized by copy number variations (CNVs), profoundly alters gene expression in cancer through direct gene dosage effects and indirect compensatory regulatory mechanisms. However, existing differential gene expression (DGE) testing methods do not differentiate between these mechanisms, conflating all expression changes, limiting biological interpretability and obscuring key genes involved in tumor progression.

To address this, we developed DeConveil, a computational framework that extends traditional DGE analysis by integrating CNV data. Using a generalized linear model with a negative binomial distribution, DeConveil models RNA-seq expression counts while accounting for copy number gene dosage effects. We proposed a more fine-grained gene decomposition into dosage-sensitive (DSGs), dosage-insensitive (DIGs), and dosage-compensated (DCGs), which explicitly de-couples changes due to CNVs and bona fide changes in transcriptional regulation. Analysis of TCGA datasets from aneuploid solid cancers resulted in notable reclassification of genes, refining and expanding upon the results from conventional methods. Functional enrichment analysis identified distinct biological roles for DSGs, DIGs, and DCGs in tumor progression, immune regulation, and cell adhesion. In a breast cancer case study, DeConveil’s CN-aware analysis facilitated the identification of both known and novel prognostic biomarkers, including long non-coding RNAs, linking gene expression signatures to survival outcomes. Utilizing these biomarkers for each gene group significantly improved patient risk stratification, yielding more accurate predictions compared to conventional methods.

These results highlight DeConveil’s ability to disentangle CNV-driven from regulatory transcriptional changes, enhancing gene classification and biomarker discovery. By improving transcriptomic analysis, DeConveil provides a powerful tool for cancer research, precision oncology, with potential applications in therapeutic target identification.

**Author Summary:** Identifying genes whose expression changes in cancer is fundamental to understand disease aetiology and to propose therapeutic targets. However, alterations to the copy number of genes, due to amplification or deletion events, can represent a significant confounder to differential expression quantification. Here we propose a simple model to correct for this confounder, identifying a finer characterization of coordinated changes in gene expression and copy number. We show on several data sets that this new characterization has prognostic value and sheds light on gene regulation in cancer.

## Introduction

Cancer is a highly heterogeneous disease characterized by extensive genomic alterations with DNA CNVs and genome aneuploidy emerging as defining hallmarks across most tumor types [1–3]. CNVs represent structural changes, such as gains or losses of specific chromosomal segments, which profoundly reshape the transcriptional landscape of cancer cells [4–6]. These changes can create gene dosage effects, amplifying or reducing mRNA transcript levels for genes within the affected regions [7,8]. Such disruptions have profound consequences for tumor progression, driving tumorigenesis [9], facilitating metastasis [10], and contributing to therapy resistance [11].

However, the relationship between somatic CNVs and gene expression is complex [4,12–19]. While some genes in altered regions exhibit expression changes that correlate with CNVs, such as oncogenes in amplified regions or tumor suppressors in deleted regions [12], many others exhibit only moderate or no expression changes [14,15], suggesting the involvement of additional regulatory mechanisms. For example, Zhou et al. (2017) [13] identified CNV-driven differentially expressed genes (DEGs) in hepatocellular carcinoma, highlighting CNVs as key drivers of transcriptional dysregulation. Mohanty et al. (2021) [19] further emphasize that CNVs alone do not dictate expression changes in aneuploid cancers; instead, gene-specific regulatory dynamics and compensatory mechanisms can modulate these effects. This complexity underscores the need for advanced statistical approaches to distinguish CNV-driven expression changes from independent regulatory alterations.

DGE analysis remains fundamental for studying transcriptomic alterations in cancer, identifying key oncogenic pathways, therapeutic targets, and biomarkers [20,21]. Widely used statistical tools for DGE analysis, such as DESeq2 [22], edgeR [23], and limma [24] employ statistical models that effectively handle RNA-seq count data, assuming gene expression changes arise solely from biological or experimental factors. However, these methods do not account for CNVs, implicitly assuming that the genes have the same CNVs across samples (or no CNVs) overall. This assumption is problematic in cancer, where aneuploidy introduces widespread CNVs that can drive gene expression changes. A key limitation is the inability to determine whether observed expression changes result from CNVs or other regulatory mechanisms. This can create significant challenges for interpreting DGE results in cancer studies, potentially obscuring key biological insights and misleading conclusions. While some progress has been made in integrating CNV data into other genomic contexts, such as DNA methylation, similar advancements in transcriptomics remain limited. For example, the ABCD-DNA tool integrates CNV data to enhance the analysis of DNA methylation [25]. However, a parallel framework for transcriptomics is still lacking.

To address these limitations, we developed DeConveil, a computational framework that explicitly integrates CNV effects into DGE analysis. DeConveil extends traditional statistical models by incorporating CNV data using a generalized linear model (GLM) with a negative binomial (NB) distribution, allowing for a more detailed interpretation of gene expression changes. This approach refines gene classification by disentangling genes whose expression is primarily driven by CNVs from those regulated through other biological mechanisms.

Application of DeConveil to aneuploid cancer datasets demonstrates its broad utility and capacity to uncover shared and specific mechanisms across cancers. In a case study on breast cancer, DeConveil provided a more refined categorization of genes based on their relationship to gene CNVs, including novel long non-coding RNAs (lncRNAs). This refined classification not only facilitated the identification of potential prognostic genes but also provided a deeper understanding of their biological roles and regulatory mechanisms underlying each gene category.

## Results

### DeConveil approach

The primary goal of `DeConveil` is to account for the influence of gene CNVs on RNA transcripts counts, when analyzing gene expression differences between contrasted conditions or sample groups. Our strategy is based on a GLM, commonly adopted in RNA-seq differential expression testing [22,23]. `DeConveil` models transcript abundance using negative binomial regression that explicitly incorporates gene- and sample-specific CN dosage as a scaling factor (see Methods). This enables correction for copy-number-driven expression shifts, which are particularly relevant in cancer, as observed in lung adenocarcinoma (LUAD) samples where transcript levels are proportional to copy number (CN) states (Fig 1A).

**Fig 1.**
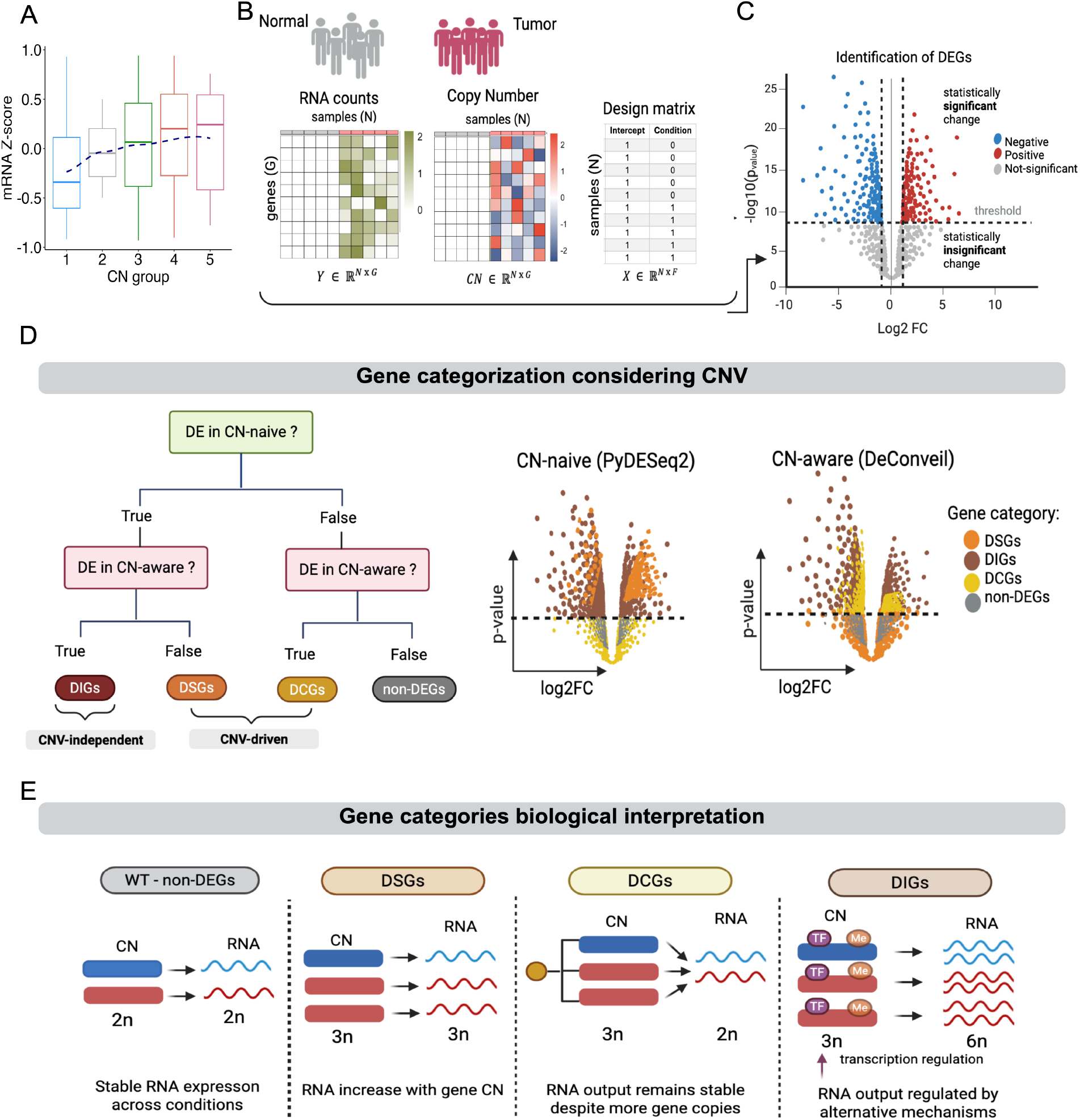
Overview of the DeConveil framework. **A**. Relationship between gene expression and DNA CNV. Boxplots show the distribution of mRNA Z-scores across five CN groups in LUAD tumor samples. The dashed blue line represents a locally weighted scatterplot smoothing (LOESS) fit, illustrating a general trend of increased expression with increasing CN. **B**. Input data and modeling design. Matched RNA-seq read counts and absolute gene CN values are provided as input matrices, with genes as rows and samples as columns. A design matrix encodes sample conditions (e.g., tumor = 1, normal = 0) and is used in our GLM framework to account for experimental groups. **C**. Statistical testing determines DEGs based on log2FC and p-values. The volcano plot highlights the criteria for selection of significantly dysregulated genes (|log2FC| > 1 and p-value < 0.05). **D**. Gene categorization framework. By comparing CN-naive (`PyDESeq2`) and CN-aware (`DeConveil`) models, genes are classified into four categories: dosage-sensitive (DSGs) are differentially expressed only in the CN-naive model; dosage-insensitive (DIGs) are differentially expressed in both models; dosage-compensated genes (DCGs) are significant only in the CN-aware model despite CN alterations; non-DEGs show no significant changes in either model. **E**. Biological interpretation of gene categories. The diagram shows how mRNA expression levels vary for different types of genes in relation to gene CN. While DSGs follow a dosage-dependent expression pattern, DIGs exhibit regulatory divergence independent of CN. DCGs show signs of compensation buffering to CN effects, and non-DEGs display stable expression across conditions.

The `DeConveil` framework integrates three key data layers (Fig 1B): RNA-seq read counts, absolute CN profiles, and a sample specific design matrix encoding experimental conditions. These inputs are used in our statistical framework to perform the log2 fold change (log2FC) and p-values calculations to perform gene classification. Genes are first classified as differentially expressed (DE) if they meet both statistical significance and effect size thresholds: adjusted p-value < 0.05 and |log2FC| > 1 (Fig. 1C). Genes that do not meet these criteria are labeled as non-differentially expressed (non-DEGs).

To disentangle regulatory-driven expression changes from those driven by CN dosage, `DeConveil` compares outputs from a standard CN-naive model (`PyDESeq2` [26]) and its CN-aware (`DeConveil`) counterpart. Based on this comparison, genes are classified into four biologically interpretable categories (Fig 1D and Fig 1E):

1. Dosage-sensitive genes (DSGs): these show RNA expression levels that scale proportionally with gene CN, consistent with gene dosage principles. While this linear assumption simplifies modeling, it may not capture all transcriptional complexities.
2. Dosage-insensitive genes (DIGs): these exhibit differential expressions that cannot be explained by CN changes alone. Instead, their expression shifts are likely driven by regulatory mechanisms such as transcription factor activity, epigenetic alterations, or post-transcriptional control [19].
3. Dosage-compensated genes (DCGs): in these genes the changes in gene CN do not linearly affect gene expression. This occurs because cells employ regulatory mechanisms to buffer these changes to maintain transcriptional homeostasis. Recent studies in cancer have demonstrated the presence of dosage compensation [14, 27–31]. For instance, when CN increases, cells may reduce transcription from amplified genes to prevent overexpression. Conversely, when CN decreases, cells may upregulate transcription to compensate for gene loss.
4. Non-DEGs: genes showing no statistically significant RNA level changes, likely reflecting transcriptional stability under the tested conditions.

This classification scheme enables more refined biological interpretation of expression changes.

The computational framework is implemented in Python as an extension of the DESeq2/PyDESeq2 statistical pipeline (https://github.com/caravagnalab/DeConveil).

### DeConveil validation using simulated data

We assessed `DeConveil’s` classification performance in distinguishing CNV-independent from CNV-driven expression changes under controlled conditions. Simulations were carried out using synthetic mRNA-seq and CNV data across varying sample sizes and gene counts (see Methods: Simulation benchmarking). Gene categories were defined to reflect distinct expression-CNV relationships (Fig 2A): DSGs, DIGs and non-DEGs. DIGs, true CNV-independent genes, serve as the true positives in our validation framework. DSGs, treated as false positives, as they should ideally be separated from the DIG category.

**Fig 2.**
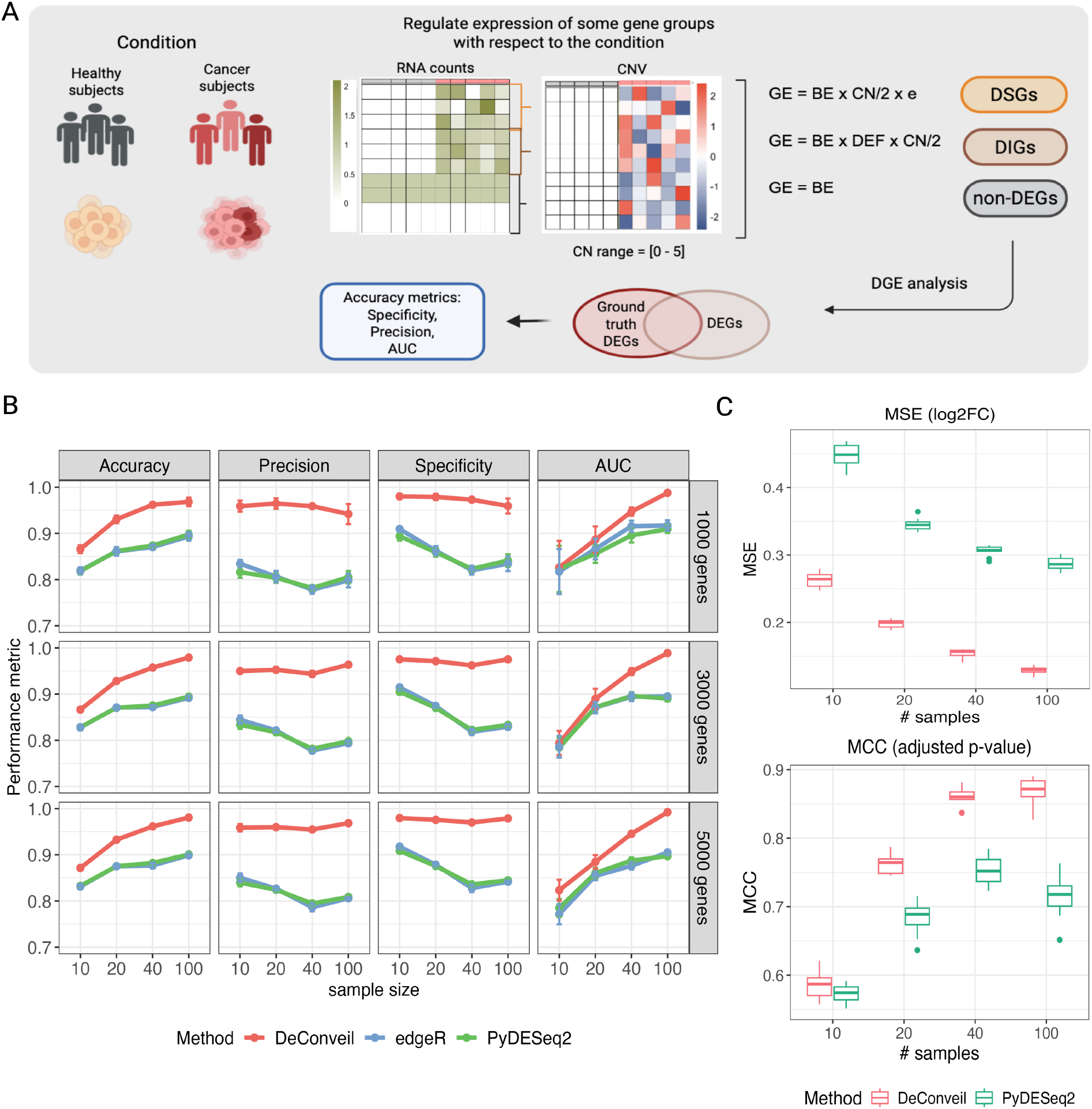
Results on simulated data. **A**. Schematic overview of the simulation setup for DGE analysis. GE is modeled as a product of baseline expression (BE) factor, CN changes, differential expression factor (DEF), and a residual error term. **B**. Performance evaluation of the `DeConveil` method on synthetic datasets with varying sample sizes (10, 20, 40, 100) and gene set sizes (1000, 3000, and 5000). Three key performance classification metrics - accuracy, precision, specificity, and AUC - were assessed and compared against CNV-naïve methods (`PyDESeq2` and `edgeR`). **C**. Evaluation of `DeConveil’s` ability to remove CNV as a confounder in effect size and significance estimation. Left: Mean Square Error (MSE) between estimated and true log2FC. Right: Matthews Correlation Coefficient (MCC) computed on adjusted p-values, assessing classification of truly DEGs.

Gene classification performance was benchmarked against CN-naive methods (`PyDESeq2` and `edgeR`), with a specific focus on identifying DIGs. It is important to highlight that `DeConveil` demonstrates superior performance in this classification task, particularly because CN-naive methods are not designed to handle such fine-grained classification. Specifically, `DeConveil` achieved higher scores in accuracy (0.86–0.98 vs. 0.82–0.90), precision (0.95–0.97 vs. 0.80–0.82), specificity (0.95–0.98 vs. 0.82–0.90), and AUROC (0.82– 0.98 vs. 0.78–0.90). (Fig 2B). This performance gap was even more pronounced in larger datasets.

To assess the impact of CN-awareness on effect size estimation, we compared each method’s estimated log2FC values to the known ground truth values used during simulation (Fig 2C). `DeConveil` achieved lower mean squared error (MSE) across all sample sizes, indicating more accurate effect size estimation. Additionally, we assessed gene detection performance using Matthew’s correlation coefficient (MCC) [32] based on adjusted p-values (Fig 2C). These results further support `DeConveil’s` superior classification accuracy compared to CN-naive methods.

To test `DeConveil’s` robustness against CNV input uncertainty, we introduced increasing levels of noise (10–25%) to the CN matrix entries and tested performance across different sample sizes (10–60). We used three metrics to assess stability (Fig S2): mean Jaccard index to measure consistency in gene group assignments, Pearson correlation (R^2^) for log2FC estimates, and Spearman correlation (R^2^) for adjusted p-value rankings. Overall, `DeConveil` demonstrated strong robustness to CN noise, particularly for DIG and non-DEG groups. These groups maintained stable classification and accurate effect size estimates across all conditions (Jaccard index > 0.85, R^2^ > 0.75), even at higher noise levels and larger sample sizes. In contrast, CN-sensitive groups (DSG and DCG), which are more dependent on accurate CN information, exhibited moderate declines in performance metrics under noise. This is particularly evident in Jaccard index and Spearman correlation MCC metric.

These results demonstrate that by incorporating CNV information, `DeConveil` not only enables more reliable classification of DEGs, but also mitigates the confounding effects of CN variation in bulk RNA-seq analysis.

### Application of DeConveil to DGE analysis

We used `DeConveil` to understand how CN corrections of gene expression influence DGE analysis results outcomes in a real scenario. We focused on evaluating the `DeConveil` ability to categorize gene expression based on our hypothesis of transcriptional effects driven by CNV. For this analysis, we selected different solid cancer types with high variability in gene CN based on availability of matched normal-tumor samples.

Using a set of 45 tumor-normal matched samples of lung adenocarcinoma (LUAD), we compared `PyDESeq2` and `DeConveil` approaches to demonstrate how our CN-informed approach enables more refined gene expression categorization by distinguishing active expression changes from passive effects caused by CNVs. To enhance the reliability of the results, we excluded genes with low expression in normal tissue (mean expression < 10 reads). After filtering, our analysis focused on 19,830 genes.

Our results reveal that CN gains and amplifications (CN 3–5) have a significant impact on gene expression, affecting approximately 75 % of DEGs (Fig 3A-C). `DeConveil` identified 875 (14.9%) DSGs, 3969 (67.6%) DIGs, and 1029 (17.5%) DCGs (Fig 3B). We then assessed the differences between `PyDESeq2` and `DeConveil` analyses by comparing the effect sizes (log2FC) and false discovery rate (FDR) across both methods (Fig 3D). For genes with neutral CNs (no CN change) used as controls, we observed concordance between the two methods (diagonal trend). However, genes affected by CN gains and amplifications exhibited higher deviations from the diagonal, confirming the influence of CN adjustment in regions with aneuploidy. Notably`, DeConveil` analysis increased FDR for amplified and gain-affected genes (Fig 3D). On average, FDRs shifted by 0.0033, with 40.75% (1837 genes) showing increased p-values, reinforcing this trend. Furthermore, 122 genes lost statistical significance (p < 0.05 in CN-naive but p > 0.05 in CN-aware analysis).

**Fig 3.**
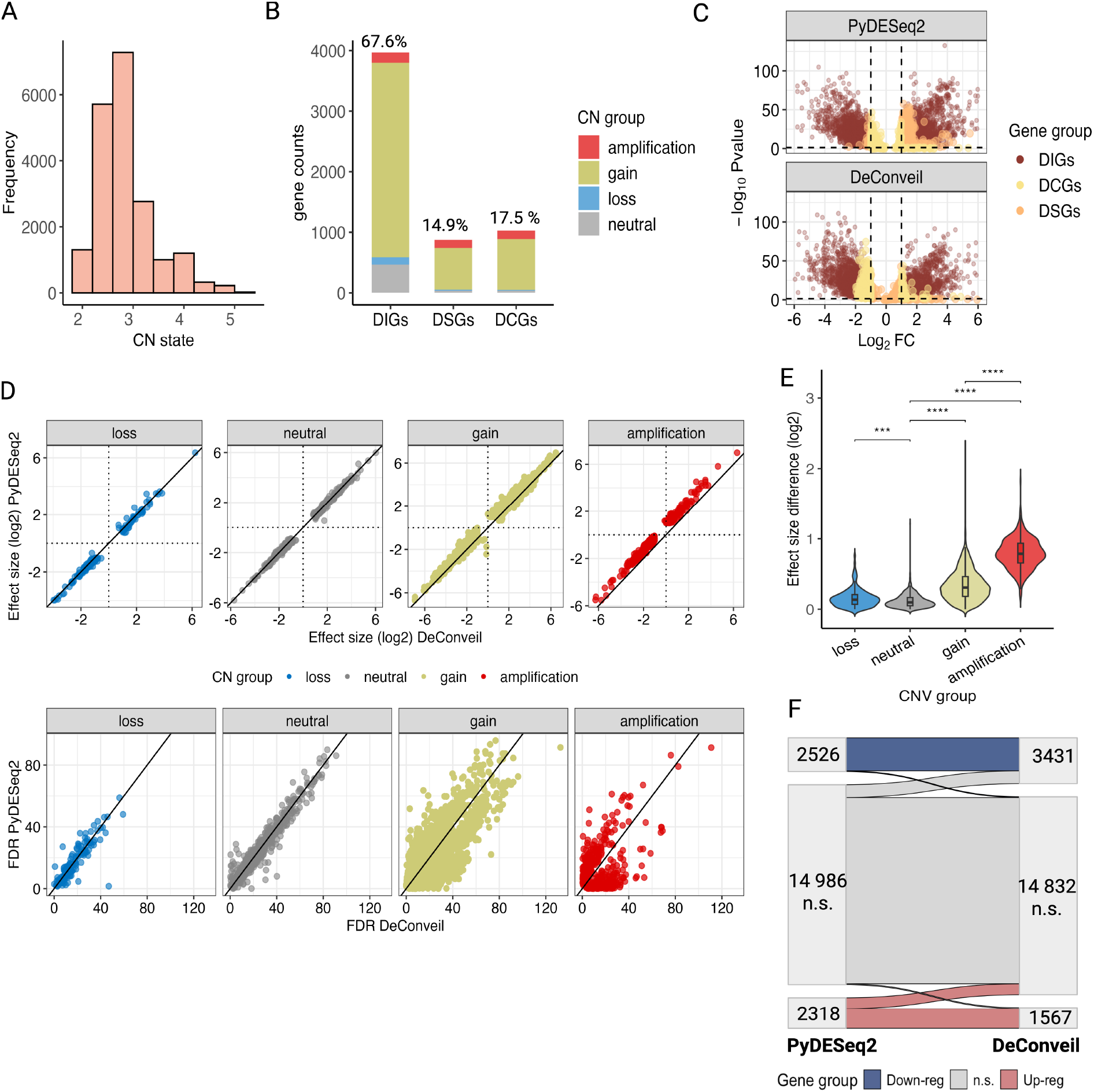
Impact of CN corrections on DGE analysis in aneuploid lung adenocarcinoma (LUAD). **A**. Histogram showing the distribution of CNV states across LUAD tumor samples. **B**. Categorization of genes based on their CNV status and classification into different gene groups: DIGs, DSGs, and DCGs. The stacked bar plot shows the proportion of each gene category, with CN states (amplification, gain, loss, and neutral). Percentages on top indicate the proportion of each group among the total gene set. **C**. Volcano plots comparing gene expression changes between the `PyDESeq2` and `DeConveil` DGE analysis approaches. Each dot represents a gene, with its position determined by logFC and p-value. Threshold for significant differential expression: |log_2_FC| > 1 and FDR < 0.05. **D**. Comparison of effect size (log2FC) and FDR (bottom row) between `PyDESeq2` and `DeConveil` models across different CN groups: loss, neutral, gain, and amplification. The diagonal reference line represents a one-to-one correlation; deviations from this line indicate differences in effect size or FDR between the two approaches. **E**. Violin plot showing the distribution of effect size differences (log_2_ scale) across CNV groups. **F**. Sankey diagram illustrating the transition of genes between expression categories in `PyDESeq2` vs. `DeConveil` analyses. Genes are grouped into upregulated (red), downregulated (blue), and non-significant (gray, n.s.) categories based on |log2FC| > 1 and FDR < 0.05. The diagram highlights shift in gene classification when `DeConveil` CN adjustments are applied.

To evaluate expression variability, we analyzed effect size differences between the methods (Fig 3E). Amplified and gain genes showed the largest effect size recalibrations of 0.79 ± 0.28 and 0.34 ± 0.26 respectively, while neutral and loss categories exhibited minimal changes (0.11 ± 0.12 and 0.14 ± 0.14) indicating less CN-driven effect. For instance, the number of downregulated genes increases after CN-aware adjustment (from 2526 to 3431), while the number of upregulated genes decreases (from 2318 to 1567), highlighting DeConveil’s ability to separate active regulatory effects from passive CNV-driven changes (Fig 3F).

We extended this analysis to four other cancer types with varying levels of CN variability (Fig S3): lung squamous cell carcinoma (LUSC), breast invasive carcinoma (BRCA), liver hepatocellular carcinoma (LIHC), and kidney renal clear cell carcinoma (KIRC). Cancers with higher CN variability (CN 1-5), such as LUSC, BRCA, and LIHC, showed a greater impact of CN corrections, as evidenced by a larger proportion of DSGs (13.7 - 21.4%) and DCGs (10.7 - 20.3%) (Fig S3, A-C). In contrast, KIRC, which exhibits lower CN variability (CN 1–3), underwent minimal gene classification shifts, with fewer DSGs (10.4%) and DCGs (8.0%). This suggests that the influence of CN corrections is less pronounced in cancers with low CN variability.

### Insights into DSGs, DIGs, and DCGs in aneuploid cancers

We further analyzed DSGs, DIGs, and DCGs identified by `DeConveil` across three epithelial origin aneuploid cancers, LUAD, LUSC, and BRCA (Fig 4A), to explore shared and private gene expression patterns and uncover functional pathways associated with each gene category.

**Fig 4.**
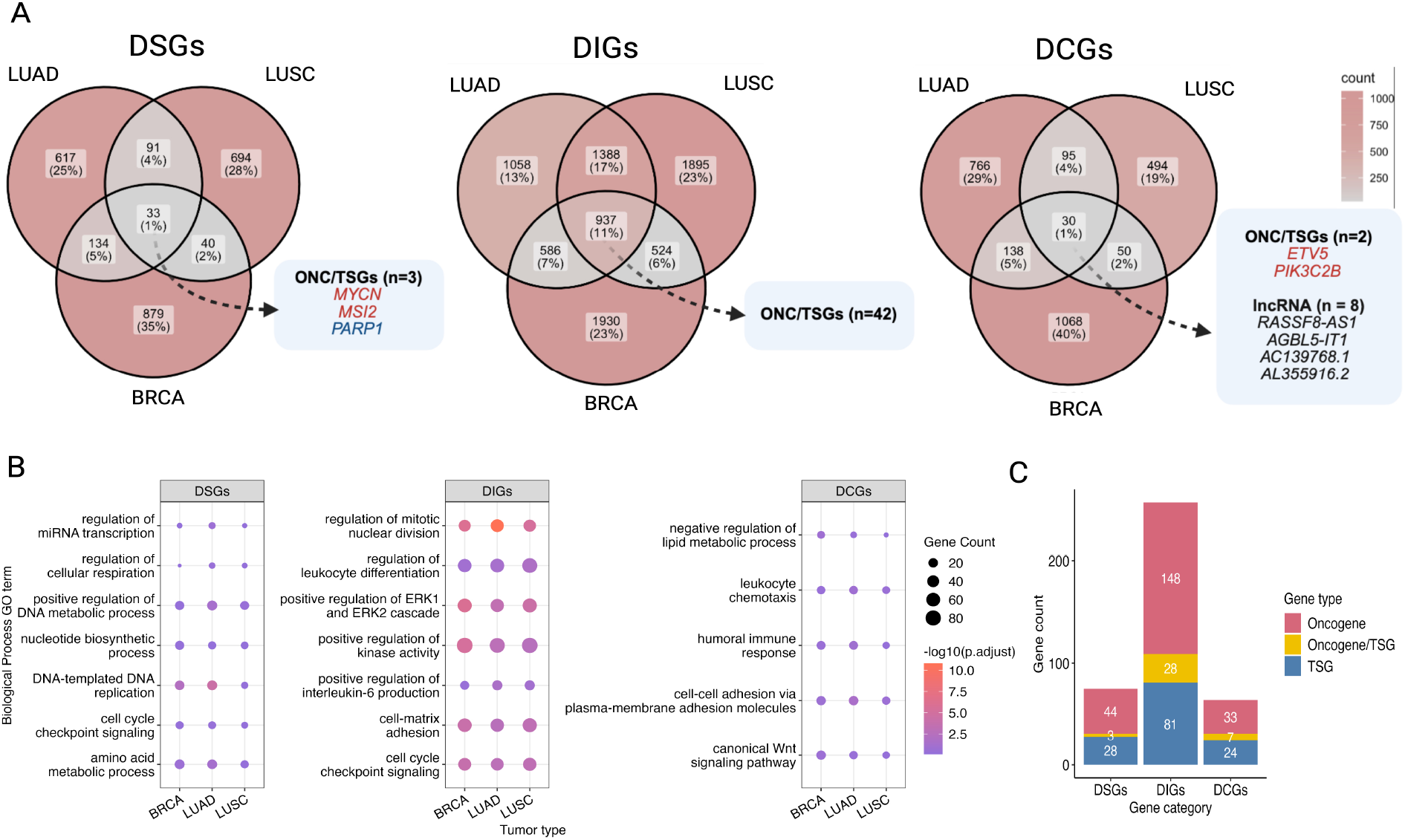
Cross-cancer comparison of DEG categories and their functional associations. Venn diagrams illustrate the overlap of DE gene categories (DSGs, DIGs, and DCGs) across three cancer types: LUAD, LUSC, and BRCA. Highlighted oncogenes (ONC) and tumor suppressor genes (TSGs) found in each category are listed in blue boxes. Additionally, 8 long non-coding RNAs (lncRNAs) were identified within the DCGs category. Genes classified differently in different cancer types are assigned to all relevant categories; thus, gene sets in this figure are not mutually exclusive. **B**. Gene Ontology (GO) term over-representation analysis for biological processes associated with DSGs, DIGs, and DCGs across LUAD, LUSC, and BRCA. The dot plots represent significantly enriched biological processes for each gene category. The size of the dots corresponds to the number of genes associated with the process, while the color represents the statistical significance of enrichment (-log_10_ adjusted p-value). **C**. Distribution of ONC and TSGs within each gene category across private DEGs of three cancer types (LUAD, LUSC and BRCA).

Among the DSGs, only 33 (0.53 %) were shared across all three cancers, including key oncogenes (*MYCN* [33], *MSI2* [34]), and tumor suppressors (*PARP1* [35]), reported to be critical for tumor progression and survival. Most DSGs are cancer specific (S2 Table) across all analyzed tumor types (11% of genes are private of the total 15%).

DIGs, more abundant than DSGs, showed broader conservation across three cancers (S3 Table), with 937 (15.2% of the total 68.6%) shared DIGs, including 42 known oncogenes and tumor suppressors. Their greater stability suggests that DIGs represent a default expression state, largely independent of CNVs, allowing tumors to maintain essential pathways despite genomic alterations.

DCGs also showed the least overlap across cancers, with only 30 (0.5% of the total 16%) shared genes, many of which are lncRNAs. This low overlap, along with the presence of lncRNAs, suggests that DCGs may function as regulatory elements influencing oncogenes and tumor suppressors activity through epigenetic or transcriptional mechanisms.

To assess the functional relevance of these shared gene categories, we performed Gene Ontology (GO) over-representation analysis (Fig 4B). The analysis revealed that shared DSGs are enriched in metabolic and cell cycle processes, confirming their importance in cancer cell survival under dosage variability. DIGs are linked to cell cycle, immune response, and oncogenic signaling pathways, supporting their role in tumor maintenance beyond CNV effects. Meanwhile, DCGs were primarily linked to immune regulation and cell adhesion, suggesting their involvement in tumor-immune interactions.

Additionally, we examined the functional significance of private cancer-specific genes within each category. For instance, functional analysis (Fig S5) linked DIGs to immune regulation and cell proliferation in LUAD, mesenchymal differentiation in LUSC, and hormone metabolism in BRCA, emphasizing their role in sustaining tumor-specific traits. Functional enrichment of private DCGs (Fig S5) highlighted their involvement in tumor-immune interactions and metabolic adaptation, including MHC complex and T-cell activation in LUAD, cytokine regulation in LUSC, and insulin secretion in BRCA.

To further refine our understanding of the significance of private genes, we mapped them to known cancer-specific oncogenes and TSGs (Fig 4C). As expected, the DIG category contained the highest number of oncogenes (n=172) and tumor suppressors (n=110), compared to DSGs (n=55 oncogenes, n=53 TSGs) and DCGs (n=53 oncogenes, n=31 TSGs). This supports the idea that DIGs encompass a broader range of critical cancer genes, many of which may operate independently of CNV effects, relying instead on regulatory mechanisms for tumor progression.

### Identification of prognostic biomarkers in breast cancer using DeConveil

In this case study, we investigated the prognostic potential of DSGs, DIGs, and DCGs identified by `DeConveil` in Breast Invasive Carcinoma (BRCA) dataset, a solid highly aneuploid tumor. From 110 paired tumor-normal samples, differential expression analysis identified 1086 (17.1%) DSGs, 3977 (62.6%) DIGs, and 1286 (20.3%) DCGs among 22 076 analyzed genes (Fig S3, B). Notably, most of these genes were primarily influenced by CN gains and amplifications.

To assess their prognostic relevance, we applied Cox proportional hazards model [36] to estimate hazard ratios (HRs) and confidence intervals (CIs). This analysis identified 142 DSGs, 161 DIGs, and 69 DCGs significantly associated with overall survival (p-value < 0.05). Further feature selection using LASSO regression highlighted key genes with prognostic potential. Several key genes emerged as significant biomarkers having prognostic potential (Fig 5A). For example, DSGs like *FBXO45* and *RAD21*, along with DIGs such as *AC113368*.*1, PEX5L*, and *GHR*, were associated with worse survival outcomes (HR > 1). Similarly, several DCGs, including *FGD5, FGFR1, MUC4, INSL3* displayed significant HRs. Many of these genes have well-established roles in breast cancer progression [37–40], metastasis [41,42], and therapy resistance. Interestingly, the analysis also identified lncRNAs within the DCG category with high prognostic potential, such as *AC079822*.*1, AC244153*.*1*, and *LINC01431*. lncRNA is known as novel gene regulatory molecules involved in cancer metabolic reprogramming and progression [43].

**Fig 5.**
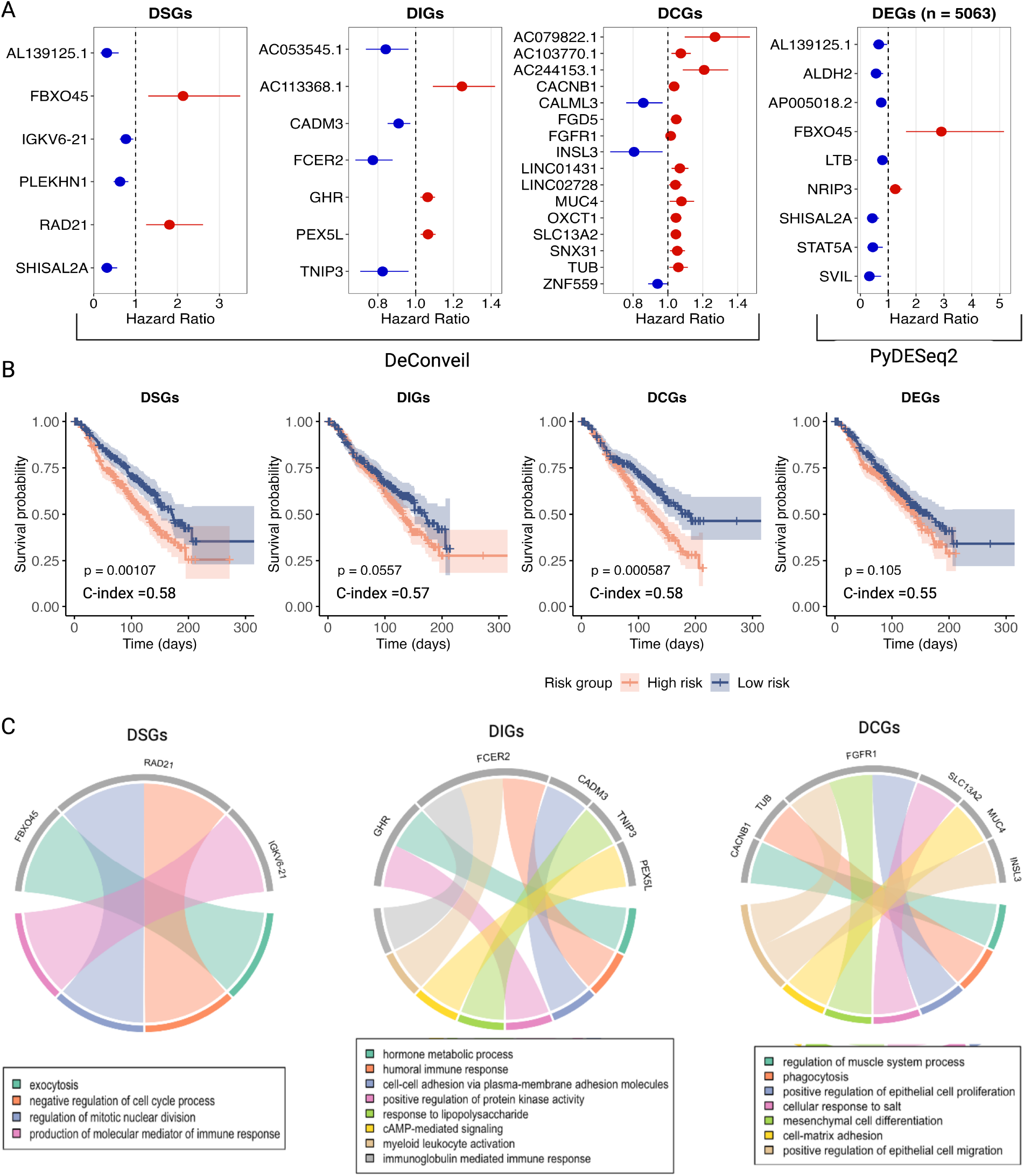
Prognostic significance of DEG categories and their functional associations. **A**. Cox proportional hazards regression analysis identifying prognostic genes within `DeConveil` DSGs, DIGs, DCGs, and PyDESeq2 DEGs. The forest plots display hazard ratios (HR) for selected prognostic genes identified using LASSO regression (p < 0.05). Genes with HR > 1 (red) indicate a high-risk association with poor survival, while those with HR < 1 (blue) are linked to better survival outcomes. **B**. Kaplan-Meier survival curves comparing high-risk (red) and low-risk (blue) patient groups based on prognostic genes identified within DSGs, DIGs, and DCGs under `DeConveil` and `PyDESeq2` models (p < 0.05). The concordance index (C-index) values indicate the predictive accuracy of the survival model. **C**. Gene Ontology (GO) enrichment analysis illustrating biological pathways associated with `DeConveil` prognostic genes in each gene category.

A comparison between `DeConveil` and `PyDESeq2` results (Fig 5A) further highlighted the advantages of DeConveil’s classification. The DeConveil identified a larger set of prognostic genes with moderate HRs, detecting key genes that `PyDESeq2` overlooked. For example, among the genes with HR > 1, only one (*FBXO45*) was identified by both approaches. This demonstrates the increased sensitivity of the `DeConveil` CN-aware approach in identifying clinically relevant genes.

To assess the predictive utility of gene signatures from each category, we calculated prognostic scores and stratified an independent cohort of 520 breast cancer patients from the METABRIC dataset into high- and low-risk groups based on median scores. Kaplan-Meier survival analysis showed that DSGs and DCGs provided the most significant separation between risk groups, with p-values < 0.001 and < 0.0006, respectively (Fig 5B). DIGs also showed a clear separation between high- and low-risk groups, but the effect was less pronounced (p < 0.05). These results highlight the high prognostic value of `DeConveil’s` gene classification framework.

Pathway enrichment analysis of the identified gene signatures (Fig 5C, Fig S6) further supported their functional distinctions. DSGs and DCGs were enriched in key cancer hallmarks, including cell cycle regulation, proliferation, adhesion, migration, and mesenchymal cell differentiation. In contrast, the DIG signature was enriched in a broader range of cancer-specific processes, such as hormone metabolism, immune response, cell adhesion, and signaling. These functional distinctions help explain the prognostic significance of these gene signatures.

## Discussion

Cancer transcriptomic heterogeneity is shaped by the interplay between genomic alterations and transcriptional regulation. Traditional DGE analyses identify expression changes across conditions but fail to assess the effect of CNVs in explaining such variations. In the context of cancers, where CNVs are prevalent, this makes it hard to distinguish which regulatory processes are impacted by CNVs, potentially obscuring key biological mechanisms underlying tumour progression.

DeConveil extends canonical approaches for DGE with CNV data, allowing a better accounting of the role of CNVs in impacting gene expression. An immediate byproduct of this new approach is the possibility of classifying genes based on the interplay among transcriptional status and CNVs. This innovative classification reveals which genes are compensated by the CNVs and those sensitive or insensitive to aneuploidy. These hypothesized biological mechanisms suggest complex regulatory mechanisms that might back up changes in gene multiplicity, underlying a complicated post-transcriptional regulatory network that acts to disrupt tissue homeostasis in the context of cancer. The implications of this new approach are also statistical. Traditional CN-naive approaches often fail to report dosage-compensated genes, suggesting that our approach might improve even established analyses of large cohorts [45–47].

We used DeConveil to analyse TCGA datasets, identifying shared and cancer-type-specific patterns of DEGs. In breast cancer, we identified known and novel prognostic biomarkers, including previously uncharacterised lncRNAs. By stratifying these genes based on CNV-related expression mechanisms, the framework improved risk group stratification and outperformed traditional DGE methods in predicting survival outcomes. This refined prognostic approach could aid in personalised treatment planning by identifying high-risk patients who may benefit from targeted interventions.

While the main contribution of our approach is to include CNVs in the DEGs picture, DeConveil still has some limitations. For instance, we assumed a linear modelling framework that may not fully capture nonlinear relationships between CNVs and gene expression. These are particularly important for high-ploidy states and RNA buffering effects that saturate the signal [4,14]. In this direction, models with diminishing returns might be considered to improve our approach.

Another key limitation is that DeConveil models CNV effects at the individual gene level without accounting for indirect transcriptional effects. CNVs in transcription factors (TFs) or signaling genes can alter the expression of downstream genes, lacking CNVs [48]. Future extensions could integrate gene regulatory network (GRN) models [49–50] to distinguish direct from indirect CNV effects.

Finally, DeConveil does not explicitly model intra-tumor heterogeneity, assuming a single CN state per gene per sample derived from segmented bulk data. As a result, subclonal CNV variation and its potential impact on gene expression are averaged out. Future versions could integrate single-cell data to improve resolution.

## Materials and Methods

### CNV-aware differential gene expression modeling

Bulk RNA-seq experiments generate a matrix of read counts **Y** ∈ ℝ^*N* × *G*^ that reflect the abundance of transcripts detected across *N* samples. For each gene *g* entry, we model the corresponding count vector ***y***_***g***_ using a Negative Binomial distribution [51–52] to account for overdispersion:

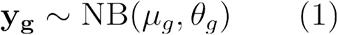

Where **y**_**g**_ represents the observed read counts for gene *g* across samples, *μ*_*g*_ is the expected gene-specific mean expression, *θ*_*g*_ captures variability in expression.

Therefore, the likelihood *L* of observed read counts for gene *g* is defined as

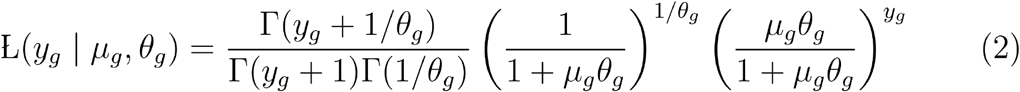

To model gene expression as a function of covariates, we use a GLM with logarithmic link:

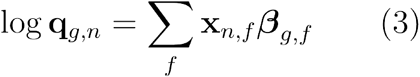

Where **q**_*g,n*_ is the expected expression level, **x**_*n,f*_ represents covariates (e.g., tumor/normal condition), **β**_*g,f*_ are the regression coefficients to be estimated.

We assume that DNA copy number measurements directly influence RNA-seq read counts. For example, a CN of 4 could result in doubled expression compared to a diploid control.

Therefore, we modify the baseline GLM to incorporate CNV effects

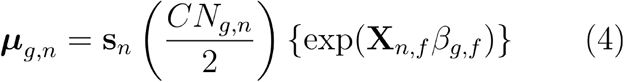

where **s**_*n*_ is a sample-specific normalization factor calculated using median-of-ratios method used in DESeq2 [22,26], while 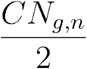 represents a gene and sample specific vector of CN dosage scaling factors. The division by 2 likely normalizes CN values relative to the diploid state (where CN = 2 is the reference). If *CN*_*g,n*_ = 2, this term becomes 1, meaning the expression is unaltered; if *CN*_*g,n*_ > 2 (amplifications), this term scales expression up, if *CN*_*g,n*_ < 2 (deletions), this term scales expression down. These components adjust the expected mean to account for systematic variables, including CN and sequencing depth.

`DeConveil` requires three input matrices: matched mRNA read counts ***Y*** ∈ ℝ^*N* × *G*^ and absolute copy number values *CN* ∈ ℝ^*N* × *G*^ (for normal diploid samples we assign CN = 2) with rows corresponding to genes and columns to samples, and design matrix ***X*** ℝ^*N* × *F*^ encoding sample conditions.

The design matrix is structured as follows: rows correspond to individual samples, and each column represents a feature. In its simplest form, **X** consists of: intercept column (constant 1 for all samples, modeling baseline expression), condition column (binary indicator: 0 = normal, 1 = tumor). For a dataset with *n* samples and *f* covariates, is an *n* × *f* matrix. The model learns an *-*dimensional coefficient vector, where *f* is the number of covariates.

`DeConveil` fits GLM for each gene and employs an empirical Bayes approach, as in DESeq2 [22,25]. Initially, the maximum likelihood estimation (MLE) is used to learn ***θ***_*g*_ and ***β***_*g*_ by maximizing log-likelihood of NB distribution. Subsequently, maximum a posteriori (MAP) estimator, namely approximate posterior estimation for GLM (apeglm) [53] method is used to apply shrinkage to both coefficients. Log2 fold change calculation is derived from estimated coefficients ***β***_*g*._

The regression coefficients ***β***_*g*_ for each gene are analyzed using the Wald test [54]. Wald test was applied to test for the statistical significance (p-value) in observed expression differences between tumor and normal sample groups. We evaluate the null hypothesis H0 : **c***β*_*g*_ = 0,where **c** is an **f**-dimensional contrast vector, that selects specific linear combinations of coefficients to test for DE. When testing multiple genes simultaneously, we adjust the resulting p-values with the Benjamini-Hochberg method [55] to reduce false positives that arise from multiple comparisons. Therefore, the results provide the log2FC calculations (magnitude of DE) and p-value (statistical significance), which are used for gene classification.

### Differential Expression classification framework

Each gene is evaluated using two DE models:

1. a CN-naive model (`PyDESeq2`), which does not account for CN effects;
2. a CN-aware model (`DeConveil)`, which incorporates sample- and gene-specific CN in the DE analysis.

A gene *g* in sample *n* is classified as DE if it meets both of the following criteria in a given model: adjusted p-value < 0.05 and absolute log_2_FC > 1. This binary DE status (1 = DE, 0 = not-DE) is evaluated independently for each gene under both models, resulting in a two-bit DE status vector per gene:

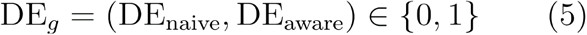

Based on DE status across both models, genes are classified into four mechanistic categories:

1. DSGs (1,0): genes that appear DE only in the CN-naive model.
2. DIGs (1,1): genes that are DE in both CN-aware and CN-naive models.
3. DCGs (0,1): genes that are DE only in the CN-aware model.
4. Non-DEGs (0,0): genes that are not DE in either model.

This classification provides a binary framework that helps to disentangle CN-driven transcriptional changes from regulatory expression shifts.

### Real datasets

The data analyzed in this study were sourced from GDC (Genomic Data Commons) Data Portal (https://portal.gdc.cancer.gov/). Specifically, we used aneuploid solid cancer datasets from TCGA, including lung adenocarcinoma (LUAD, 100 samples), breast invasive carcinoma (BRCA, 220 samples), liver hepatocellular carcinoma (LIHC, 102 samples), lung squamous cell carcinoma (LUSC, 100 samples), and kidney renal clear cell carcinoma (KIRC, 146 samples). We downloaded matched primary tumor and normal samples, including absolute gene-level copy number data, mRNA-seq read counts, and clinical information, using the TCGAbiolinks Bioconductor R package (v.2.30.4). The METABRIC DNA CN data and gene expression RNA-seq data were downloaded from the cBioPortal database (https://www.cbioportal.org/).

### Data preprocessing

In RNAseq data, genes with low expression in normal tissue were filtered out from the analysis (mean expression across samples <10 read counts) to minimize noise. The genes obtained after filtering were used in further differential expression tests. Significantly DEGs were identified based on the following criteria: | log2FC| ≥ 1 and *p* < 0.05.

For the exploratory analysis of the relationship between gene CN and mRNA expression, z-score normalization was applied to the log-transformed mRNA-seq data. For the CN data, the mean CN value for each gene was calculated across samples, and CN states were categorized as follows: 1 (CN mean > 0.0 and ≤ 1.7), 2 (CN mean > 1.7 and ≤ 2.5), 3 (CN mean > 2.5 and ≤ 3.5), 4 (CN mean > 3.5 and ≤ 4.5), and 5 (CN mean > 4.5). Additionally, Principal Component Analysis (PCA) and k-means clustering were applied to select patients for exploratory analysis based on their CN profiles.

### CNV groups definition

For the TCGA datasets used to test `DeConveil,` CNV groups were defined based on the following criteria: neutral (CN mean > 1.7 and ≤ 2.5), gain (CN mean > 2.5 and ≤ 3.5), and amplification (CN mean > 3.5). Conversely, CN loss was categorized as loss for genes exhibiting CN values 0 or 1 in at least 25% of tumor samples that were considered as frequently deleted.

### Simulation benchmarking

#### Validation of DeConveil classification performance

To simulate baseline mRNA read counts we first fitted the DESeq2 [22] model to the TCGA-BRCA real dataset to obtain empirical estimates of gene-wise mean expression and dispersion parameters. These parameters were used to generate synthetic count data using the *generateSyntheticData* function from compcodeR Bioconductor R package (v.1.38.0) [56].

We defined three gene categories to model different expression-CNV relationships:

– DSGs (CNV-driven signal genes):

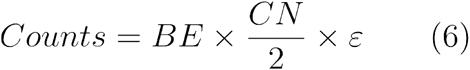
DIGs (CNV-independent genes):

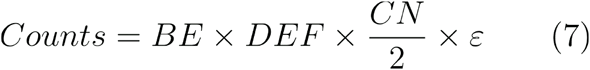
non-DEGs:

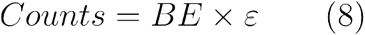

Here, *BE* represents the baseline expression level drawn from the fitted model, *CN* is the simulated copy number sampled from empirical TCGA-BRCA CNV data, *DEF* is a differential expression factor applied only to DIGs, and *ε* introduces multiplicative noise. Specifically, CN values were drawn from a realistic distribution reflecting somatic CNV profiles, with values ranging from 0 (deep deletion) to 6 (high-level amplification). Each dataset included 10% DSGs, 40% DIGs, and 50% non-DEGs to ensure a balanced test scenario [57,58].

Simulations were performed across various scenarios, varying sample size per condition (n = 10, 20, 40, 100) and varying the number of genes (n = 1000, 3000, 5000). For each simulation setting we simulated 10 independent datasets to ensure statistical robustness. `DeConvel’s` performance was compared to CN-naive statistical tools for differential expression analysis, such as `PyDESeq2` and `edgeR`. Each method’s results were evaluated against the ground truth to assess their ability to correctly classify DE genes.

We measured the ability of each method to detect DE genes using the following definitions based on a confusion matrix framework:

– True Positives (TP): correctly identified DIGs,
– False Positives (FP): DSGs or non-DEGs incorrectly identified as DE,
– True Negatives (TN): correctly identified non-DEGs.
– False Negatives (FN): DIGs missed by the method.

From these counts, we calculated:

– True Positive Rate (TPR) = TP / (TP + FN)
– False Positive Rate (FPR) = FP / (FP + TN)
– Precision = TP / (TP + FP)
– Accuracy = (TP + TN) / total genes
– Specificity = TN / (TN + FP)
– AUROC: area under the ROC curve summarizing the trade-off between TPR and FPR.

### Evaluation of DeConveil’s ability to remove CNV as a confounder

To assess how well `DeConveil` mitigates the confounding influence of CNV on effect size and significance estimates, we designed simulations focused on comparing ground truth and inferred values. We used a similar simulation strategy as in the classification performance benchmark, with the following adjustments: the total number of genes was fixed at n=1000, and a subset of 40% of genes were simulated as differentially expressed. To simulate CNV-confounding genes, we introduced signal distortion by scaling gene expression counts with a 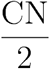 factor.

We compared `DeConveil` performance against CN-naive method `PyDESe2`. Specifically, we compared the estimated log2FC from each method to ground truth values used in the simulation. We computed the mean squared error (MSE) between estimated and true log2FC values across all genes to quantify effect size accuracy. Lower MSE values indicate more accurate and unbiased effect size estimation. Additionally, we evaluated the Matthews Correlation Coefficient (MCC) for DE gene classification, based on adjusted p-values.

### Robustness of DeConveil to CNV estimation noise

To simulate realistic uncertainty in CNV quantification we perturbed 5–20% of randomly selected CN matrix entries with additive discrete noise values. Specifically, we sampled these values from a weighted distribution: [−2 (5%), −1 (30%), 0 (30%), 1 (30%), 2 (5%)]. This weighting scheme reflects common CNV estimation errors, where moderate deviations are more frequent than extreme ones, mimicking variability typically encountered in low-purity or low-coverage tumor samples.

Both RNA-seq counts and CN profiles were simulated by subsampling from real aneuploid TCGA cancer samples, ensuring biologically plausible input patterns.

To quantify robustness, we evaluated:

1. Gene classification stability using the Jaccard index [59], calculated between the sets of genes assigned to each group (DIGs, DSGs, DCGs, and non-DEGs) in the noise-free and noise-perturbed datasets. The index was computed as

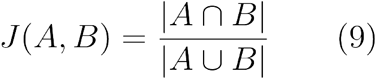

where *A* and *B* are gene group memberships before and after noise. Results were averaged across 5 replicates and noise levels. A high Jaccard index (closer to 1) indicates strong agreement and stability of classification.
2. Effect size stability via Pearson correlation (R^2^) between estimated and true log2 fold changes.
3. Significance stability using Spearman correlation (R^2^) of adjusted p-value rankings (FDR) All metrics were computed across 5 simulation replicates and sample sizes (n = 10, 20, 40, and 60).

### Functional enrichment analysis

Overrepresentation analysis of DE gene categories identified by `DeConveil` was performed using the Gene Ontology database provided by clusterProfiler (v.4.2.2) R/Bioconductor package [60]. For each gene category (DSGs, DIGs, and DCGs) we performed a hypergeometric test using the function *enrichGO*. We focused on biological process (BP) GO terms, filtering for gene sets with sizes between 10 and 350 genes. A p-value cutoff of 0.05 was used as the significance threshold for GO term identification.

### Survival analysis

We used gene groups identified by `DeConveil` to identify prognostic gene signatures associated with patient survival outcomes. Log-normalized RNA-seq expression data from each gene category (DSGs, DIGs, and DCGs) were scaled by corresponding CN values to account for gene dosage effects. These expression data were then integrated with clinical time-to-event data for survival modeling.

As a first step, we applied the Cox proportional hazards model using the survival R package (v.3.8-3) to identify genes significantly associated with overall survival. Genes with an adjusted p-value < 0.05 were retained for further analysis. To reduce dimensionality and select the most predictive features, we applied LASSO-penalized Cox regression [61] using the glmnet R package (v.4.1-8). Genes with non-zero LASSO coefficients were selected to form the final prognostic gene signature. A prognostic score was then computed for each patient as a weighted linear combination of the expression levels of these selected genes, using their corresponding LASSO-derived coefficients. Based on the median prognostic score, patients were stratified into high-risk (score > median) and low-risk (score ≤ median) groups.

To assess the survival differences between these groups, we generated Kaplan–Meier survival curves and performed log-rank tests. A p-value < 0.05 was considered statistically significant. For comparative analysis, we also evaluated DE genes identified using a CN-naive model, where gene expression was not adjusted for CN effects. These DE genes were subjected to the same survival analysis pipeline.

For external validation of the model’s predictive performance, we used the METABRIC dataset (n = 520 patients). The previously derived LASSO-selected gene signature was applied to this independent cohort, and prognostic scores were calculated accordingly. Model performance was evaluated using the concordance index (C-index), which quantifies the concordance between predicted risk scores and observed survival outcomes.

## Acknowledgments

The authors acknowledge support from the Italian Association for Cancer Research (AIRC) under grant MFAG 2020 -ID. 24913 project (PI - GC) and IG 27631 (PI - GS). GC acknowledges financial support under the National Recovery and Resilience Plan (NRRP), Mission3224, Component 2, Investment 1.1, Call for tender No. 1409 published on 14.9.2022 by the Italian Ministry of University323 and Research (MUR), funded by the European Union – NextGenerationEU– CUP J53D23015060001, as well as324 under Decreto Direttoriale No. 104 published on 02-02-2022 by MUR (NextGenerationEU– CUP J53D23003860006),325 under the PNRR-MAD-2022-12375673 (Next Generation EU, M6/C2 CALL 2022) and under the PNRR-MCNT2-3262023-12378037 (Next Generation EU, M6/C2 CALL 2023), both published by the Italian Ministry of Health, Rome,327 Italy. GS acknowledges co-funding from Next Generation EU, in the context of the National Recovery and Resilience Plan, Investment PE1 - Project FAIR “Future Artificial Intelligence Research”. This resource was co-financed by the Next Generation EU [DM 1555 del 11.10.22].

## Data availability statement

All data and code used for running experiments, model fitting, and plotting is available on a GitHub repository at https://github.com/kdavydzenka/deconveilCaseStudies. We have also used Zenodo to assign a DOI to the repository: 10.5281/zenodo.15100458.

DeConveil is available as an open-source package written in Python: https://github.com/caravagnalab/DeConveil.

