## Supplementary material for "Extending differential gene expression testing to handle genome aneuploidy in cancer": Suppl.Figures_and_Tables

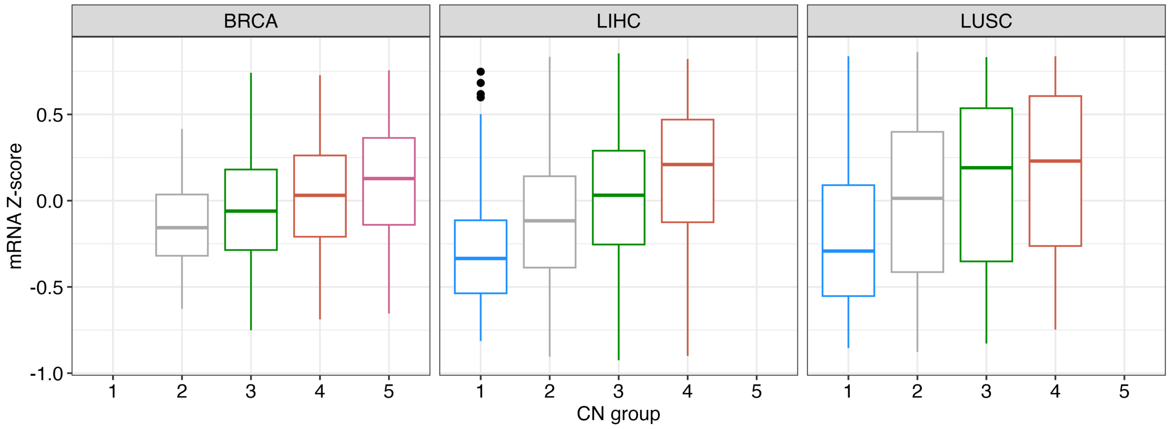


**S1 Fig. Boxplots show the relationship between mRNA Z-score and CN groups across tumor types (BRCA, LIHC, LUSC).**

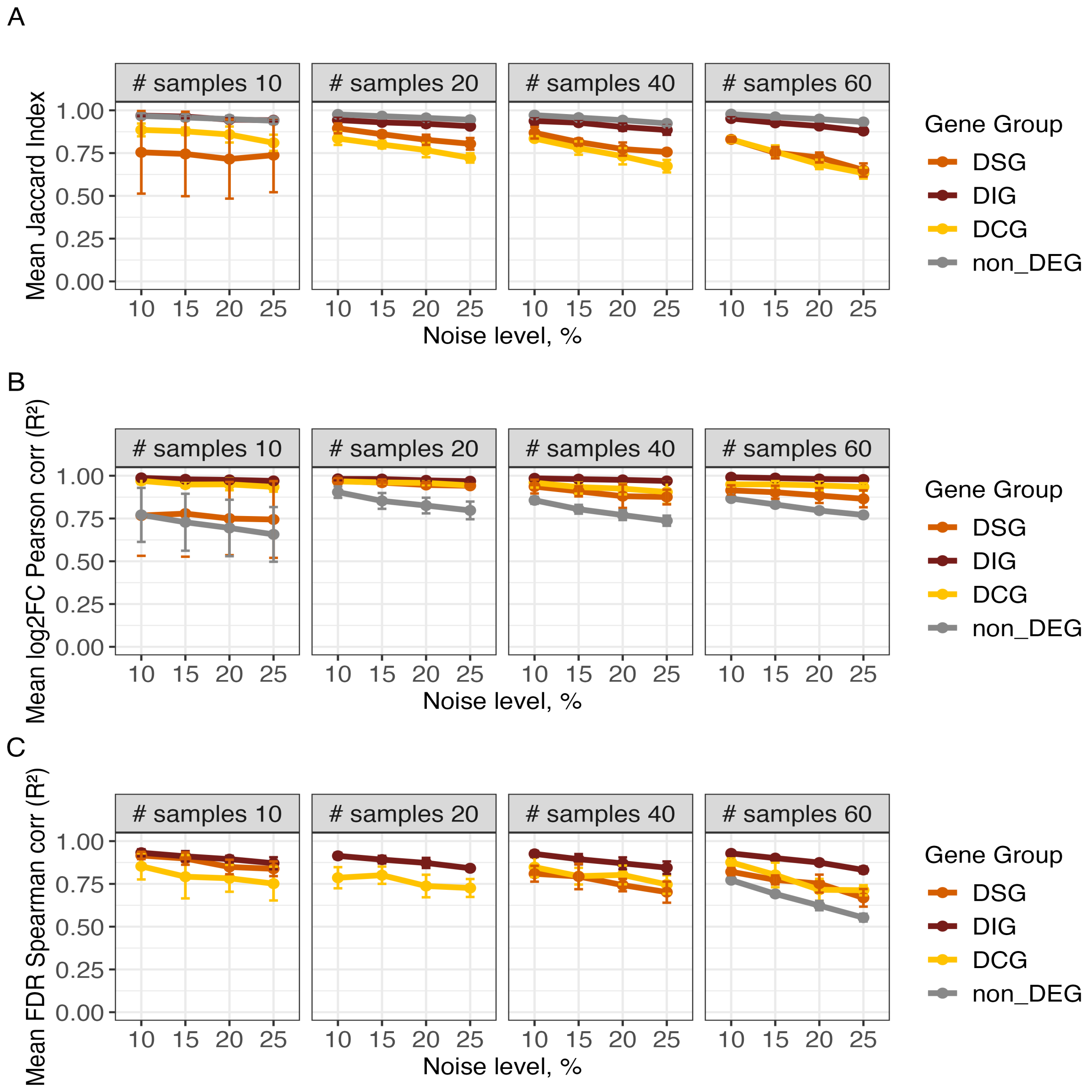


**S2 Fig. Robustness of `DeConveil` to CNV estimation noise across gene groups and sample sizes.** Simulated CN values were randomly perturbed in 5%, 10%, 15%, or 20% of the gene-sample matrix using additive discrete noise drawn from [−2, 2]. Panels show metrics across 5 replicates per condition. **A.** Mean Jaccard index across replicates assessing gene group classification stability. **B.** Pearson R2 correlation between noise-free and noisy log2FC. **C.** Spearman R2 correlation between FDR values under noise-free vs. noisy conditions. `DeConveil` is resilient to CN noise, especially in DIG and non-DEG groups, while CN-sensitive groups (DSG, DCG) show moderate declines at higher noise levels and larger sample sizes.

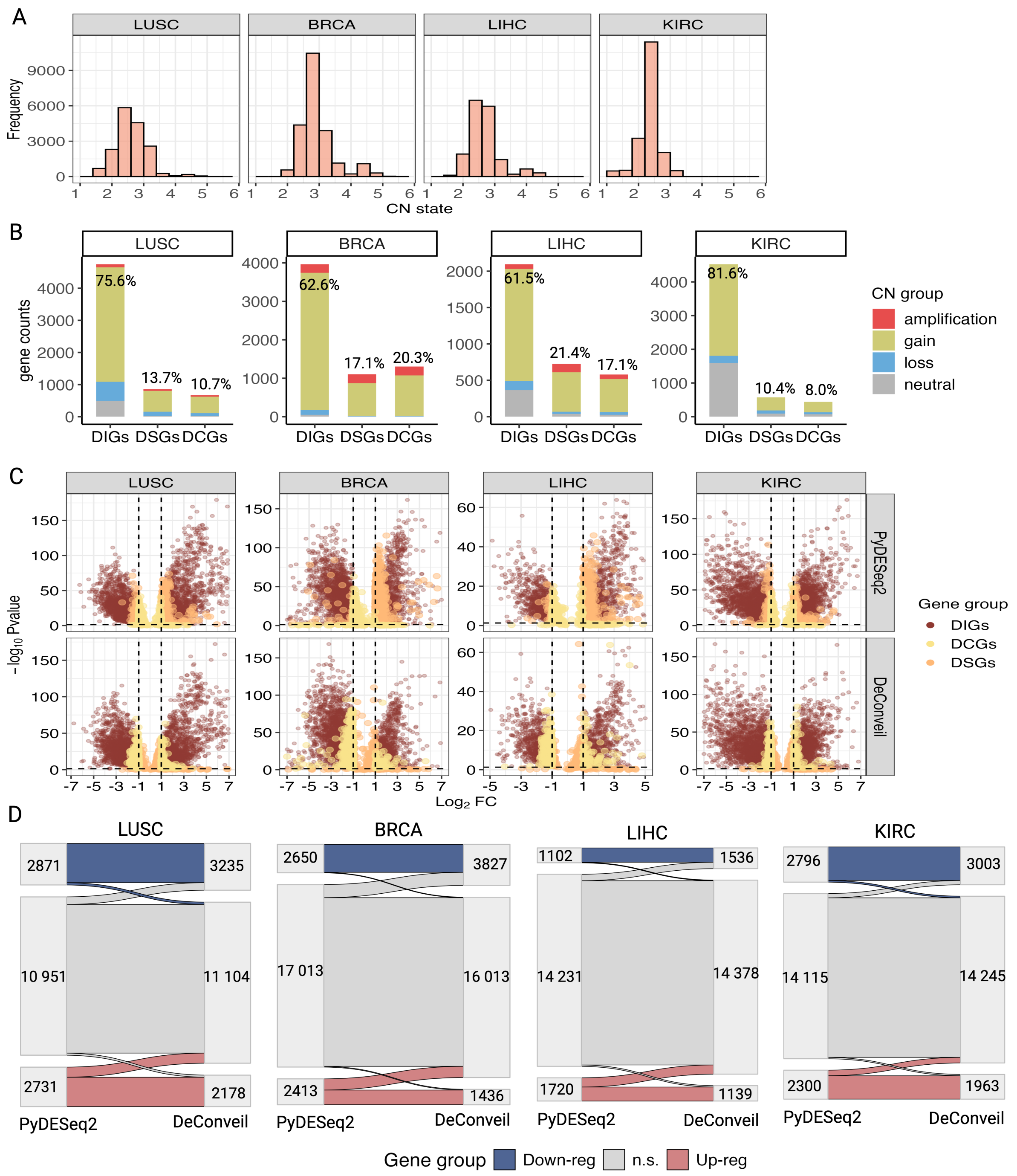

**S3 Fig. Impact of CN corrections on DGE analysis across aneuploid cancer types (LUSC, BRCA, LIHC, KIRC). A.** Histogram showing the distribution of CNV states across tumor samples. **B.** Categorization of genes based on their CNV status and classification into different gene groups: DIGs, DSGs, and DCGs. The stacked bar plot shows the proportion of each gene category, with CN states (amplification, gain, loss, and neutral). Percentages on top indicate the proportion of each group among the total gene set. **C.** Volcano plots comparing gene expression changes between the `PyDESeq2` and `DeConveil` DGE analysis approaches. Each dot represents a gene, with its position determined by log2FC and p-value. Threshold for significant DE: $|\log_2 FC| > 1$ and FDR < 0.05. **D.** Comparison of effect size (log2FC) and FDR (bottom row) between `PyDESeq2` and `DeConveil` models across different CN groups (loss, neutral, gain, and amplification). The diagonal reference line represents a one-to-one correlation; deviations from this line indicate differences in effect size or FDR between the two approaches. **E.** Violin plot showing the distribution of effect size differences ($log_2$ scale) across CNV groups. **F.** Sankey diagram illustrating the transition of genes between expression categories in `PyDESeq2` vs. `DeConveil` analyses. Genes are grouped into upregulated (red), downregulated (blue), and non-significant (gray, n.s.) categories based on $|log2FC| > 1$ and $FDR < 0.05$. The diagram highlights shift in gene classification when CN adjustments are applied.

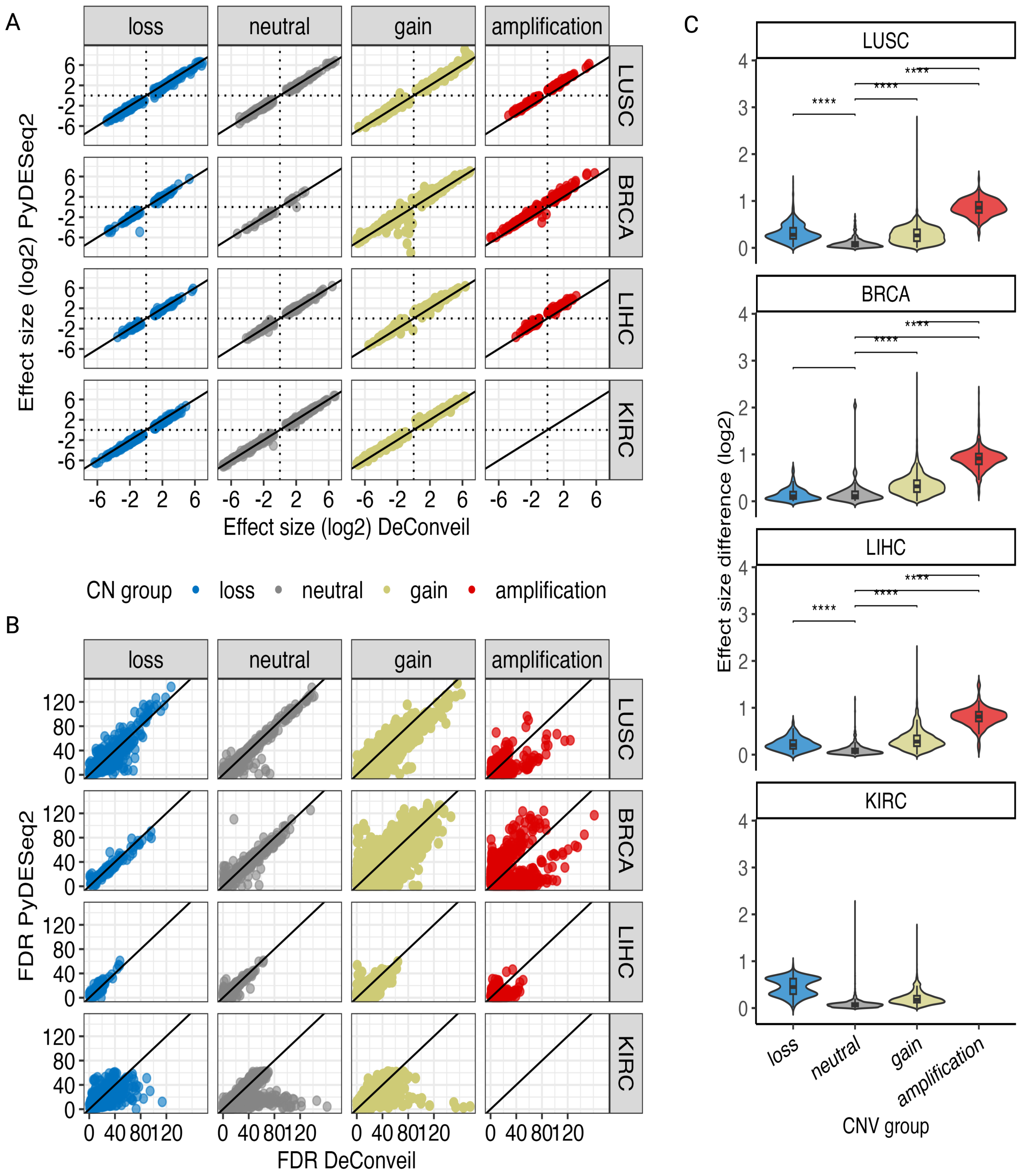


**S4 Fig. Impact of CN corrections on DGE analysis across aneuploid cancer types (LUSC, BRCA, LIHC, KIRC). A.** Comparison of effect size (log2FC) between `PyDESeq2` and `DeConveil` models across different CN groups (loss, neutral, gain, and amplification). The diagonal reference line represents a one-to-one correlation; deviations from this line indicate differences in effect size or FDR between the two approaches. **B.** Comparison of FDR between `PyDESeq2` and `DeConveil` models across different CN groups. **C.** Violin plot showing the distribution of effect size differences ($log_2$ scale) across CNV groups.

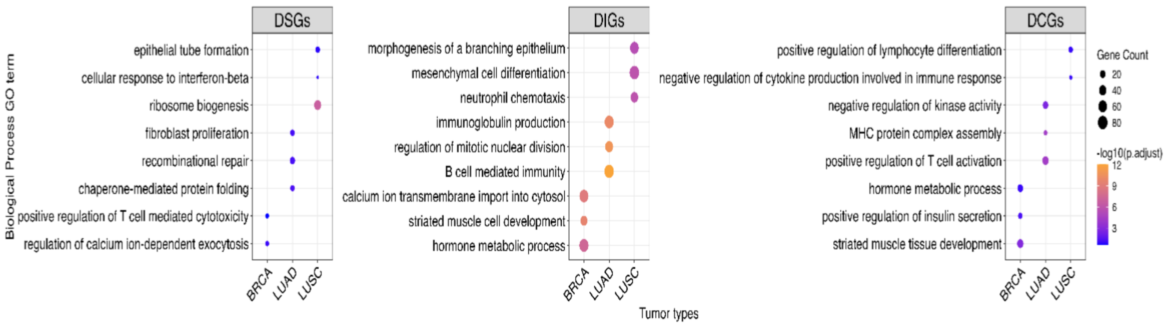


**S5 Fig. Gene Ontology (GO) term enrichment analysis for biological processes associated with DSGs, DIGs, and DCGs across LUAD, LUSC, and BRCA.** The dot plots represent significantly enriched biological processes for each gene category. The size of the dots corresponds to the number of genes associated with the process, while the color represents the statistical significance of enrichment (-$\log_{10}$ adjusted p-value).

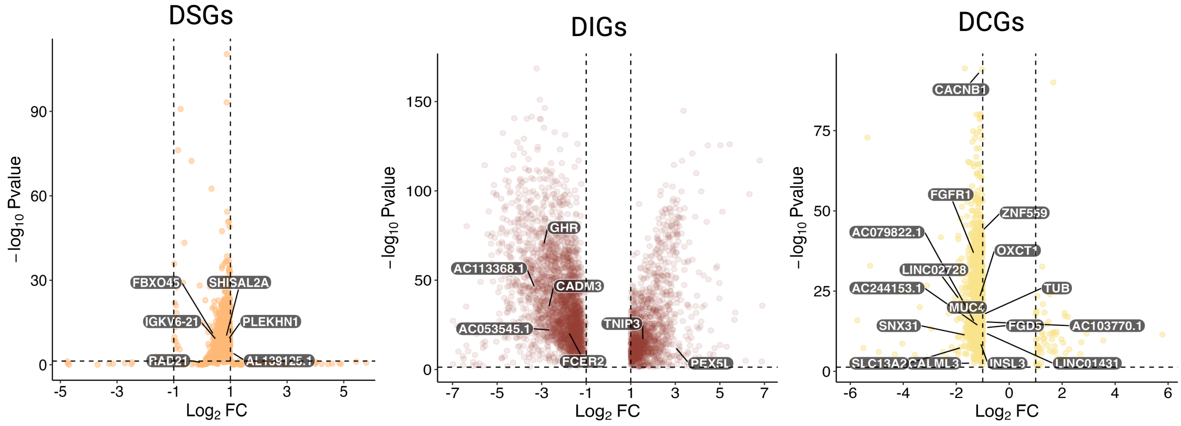


**S6 Fig. Volcano plots showing prognostic genes identified with Cox/LASSO regression within each gene category (DSGs, DIGs, DCGs).**

**S1 Table. Summary of private gene distribution (%) across gene categories (DSGs, DIGs, and DCGs) and cancer types (LUAD, LUSC, BRCA).** This table provides insights into the relative distribution of private genes within each gene category and cancer type, and their overall background proportions.

| Gene category | LUAD, % | LUSC, % | BRCA, % | proportion mean (n) | background proportion, % |
|---|---|---|---|---|---|
| **DSGs** | 10.0 | 11.2 | 14.2 | 11.8 | 15.2 |
| **DIGs** | 17.1 | 30.7 | 31.3 | 26.4 | 68.6 |
| **DCGs** | 12.4 | 8.0 | 17.3 | 15.5 | 16.1 |

**S2 Table. Summary of gene categories across three cancer types (LUAD, LUSC, BRCA).** This table provides the number of genes in three different gene categories (DSGs, DIGs, and DCGs). The table includes the mean number of genes per category, their proportion in percentage, and the number and proportion of shared genes among the categories.

| Gene category | LUAD (n) | LUSC (n) | BRCA (n) | gene mean (n) | proportion, % | shared genes (n) | proportion shared, % |
|---|---|---|---|---|---|---|---|
| **DSGs** | 875 | 858 | 1086 | 940 | 15.2 | 33 | 0.53 |
| **DIGs** | 3969 | 4744 | 3977 | 4230 | 68.6 | 937 | 15.2 |
| **DCGs** | 1029 | 669 | 1286 | 995 | 16.1 | 30 | 0.49 |
